# Discovery of paralogous GnRH and corazonin signaling systems in an invertebrate chordate

**DOI:** 10.1101/2023.03.06.531269

**Authors:** Luis Alfonso Yañez Guerra, Meet Zandawala

**Affiliations:** Living Systems Institute, University of Exeter, Exeter EX4 4QD, UK; Neurobiology and Genetics, Theodor-Boveri-Institute, Biocenter, Julius-Maximilians-University of Würzburg, Am Hubland, 97074 Würzburg, Germany

**Keywords:** Neuropeptide, GPCR, Evolution, Gonadotropin-releasing hormone, Adipokinetic hormone, Corazonin, Hidden Markov Model

## Abstract

Gonadotropin-releasing hormone (GnRH) is a key regulator of reproductive function in vertebrates. GnRH is related to the corazonin (CRZ) neuropeptide which influences metabolism and stress responses in insects. Recent evidence suggests that GnRH and CRZ are paralogous and arose by a gene duplication in a common ancestor of bilaterians. Here we report the identification and complete characterization of the GnRH and CRZ signaling systems in the amphioxus *Branchiostoma floridae*. We have identified a novel GnRH peptide (YSYSYGFAP-NH_2_) that specifically activates two GnRH receptors and a CRZ peptide (FTYTHTW-NH_2_) that activates three CRZ receptors in *B. floridae*. The latter appear to be promiscuous, as two CRZ receptors can also be activated by GnRH in the physiological range. Hence, there is a potential for cross-talk between these closely-related signaling systems. Discovery of both the GnRH and CRZ signaling systems in one of the closest-living relatives of vertebrates provides a framework to discover their roles at the transition from invertebrates to vertebrates.

**Significance:** Identifying neuropeptides and resolving the relationships of neuropeptides across different phyla is challenging due to their short sequences. Here we resolve a controversy regarding the identities and evolutionary relationships of a family of homologous neuropeptides that include the well-known human reproductive hormone gonadotropin-releasing hormone (GnRH) and insect stress hormone corazonin (CRZ). We have discovered bona fide GnRH and CRZ signaling systems in *Branchiostoma* which provides a basis for investigating the evolution of the physiological roles of these neuropeptides in one of the closest living relatives of vertebrates.

## Introduction

Neuropeptides are the most diverse class of neuronal signaling molecules. They are found throughout the animal kingdom, including in animals without a nervous system (Nassel and Zandawala 2019; Sachkova, et al. 2021; Senatore, et al. 2017; Yanez-Guerra, et al. 2022). The origins of most neuropeptide families can be traced to a common ancestor of bilaterian animals (Jekely 2013; Mirabeau and Joly 2013). Consequently, orthologs of several vertebrate neuropeptide signaling systems are found in invertebrates. One such conserved neuropeptide signaling pathway that has been widely-studied is the gonadotropin-releasing hormone (GnRH) system which regulates reproductive function in humans and other vertebrates (Tsutsumi and Webster 2009). GnRH is orthologous to the adipokinetic hormone (AKH) in invertebrates such as arthropods and regulates energy homeostasis (Bharucha, et al. 2008; Staubli, et al. 2002; Zandawala, et al. 2018). The GnRH/AKH signaling system is closely related to the corazonin (CRZ) signaling pathway which influences metabolism and stress responses in insects (Kubrak, et al. 2016; Veenstra 2009; Zandawala, et al. 2021; Zhao, et al. 2010). Recent evidence from the starfish *Asterias rubens* revealed that the GnRH and CRZ signaling systems are in fact paralogous and arose by gene duplication in a common ancestor of bilaterians (Tian, et al. 2016). This study put an end to a long-standing debate on the precise relationship between GnRH and CRZ (Zandawala, et al. 2018), exemplifying the power of deuterostomian invertebrates in resolving the evolution of bilaterian neuropeptide signaling systems. Independently, the AKH signaling system also appears to have duplicated in a common ancestor of the arthropods to give rise to the AKH/CRZ-related peptide (ACP) signaling, whose function remains elusive (Hansen, et al. 2010).

Occasionally, both the neuropeptide sequence/structure and function tend to be conserved across animals, as evident for insulin which regulates lifespan in most animals (Barbieri, et al. 2003). Hence, invertebrates such as insects and nematodes have served as good models to decipher the functions and modes-of-action of some conserved neuropeptides (Nässel and Wu 2022; Van Sinay, et al. 2017). However, in most cases, including GnRH and AKH, vertebrate and insect counterparts of the same neuropeptide family regulate different functions. This is perhaps due to the large evolutionary distance separating these species. Therefore, there is a need to establish other model systems that are more closely related to vertebrates than insects to study neuropeptide function. The amphioxus *Branchiostoma floridae* is best suited to address this need. Amphioxi are chordates and represent one of the closest living relatives of vertebrates (Putnam, et al. 2008). Discovering neuropeptide signaling systems in *B. floridae* can thus provide a framework to unravel the functions of neuropeptides in this species which occupies a unique position in animal phylogeny.

Previously, one neuropeptide precursor and four receptors belonging to the GnRH/CRZ superfamily were identified in the *B. floridae* genome (Roch, et al. 2014b; Tello and Sherwood 2009). Phylogenetic analysis revealed that two of these receptors grouped with protostomian CRZ receptors and the other two receptors grouped with GnRH/AKH receptors. A putative mature peptide (pQILCARAFTYTHTWNH_2_) encoded by this novel precursor was classified as a GnRH-like peptide despite the fact that it activated one of the two CRZ-like receptors in the high nanomolar range (Roch, et al. 2014b). Regardless of the nomenclature, it is evident that additional and/or more specific peptide ligands for the *B. floridae* GnRH and CRZ receptors remain to be discovered. Here, we report the discovery of the authentic *B. floridae* GnRH precursor using a Hidden Markov Model (HMM)-based search. We show that the putative mature peptide encoded by this precursor can activate both the GnRH receptors. We also discover an additional previously unidentified CRZ receptor and identify the putative endogenous ligands for all three CRZ receptors.

## Results and Discussion

### Identification of the authentic *B. floridae* GnRH precursor

To identify CRZ-like and GnRH-like precursors in *B. floridae*, we searched the *Branchiostoma* transcriptomes using a custom-generated HMM for GnRH/CRZ precursors. Our sensitive search uncovered a single novel precursor in *Branchiostoma belcheri* and *B. floridae*. Since the precursor previously discovered by Roch et al. (2014) encodes a peptide (Figure S1) which activates a CRZ receptor, it would be more appropriate to reclassify it as the *B. floridae* CRZ precursor (Roch, et al. 2014b). With this in mind, we hypothesized that our newly-identified precursor encodes the elusive *B. floridae* GnRH (Figure 1A and S2). Sequence analysis revealed that this precursor is predicted to generate a single copy of the peptide YSYSYGFAP-NH_2_. Interestingly, this peptide lacks the N-terminal glutamine (which gets converted to pyroglutamic acid *in vivo*) that is a conserved feature across vertebrate GnRH and arthropod AKH peptides (Figure 1B). *In silico* analyses of the *B. floridae* CRZ precursor suggests that it also encodes a single copy of the mature peptide FTYTHTW-NH_2_ (Tian, et al. 2016; Zandawala, et al. 2018). This peptide is much shorter than the putative mature peptide (pQILCARAFTYTHTW-NH_2_) predicted previously (Roch, et al. 2014b; Zandawala, et al. 2018). Sequence analysis indicates that the seven amino acid residues at the N-terminal end of the extended peptide are in fact part of the signal peptide which immediately precedes the predicted mature peptide (Figure 1A). Consequently, *B. floridae* CRZ also lacks the highly-conserved N-terminal glutamine and is the shortest CRZ discovered to date (Figure 1C).

**Figure 1:**
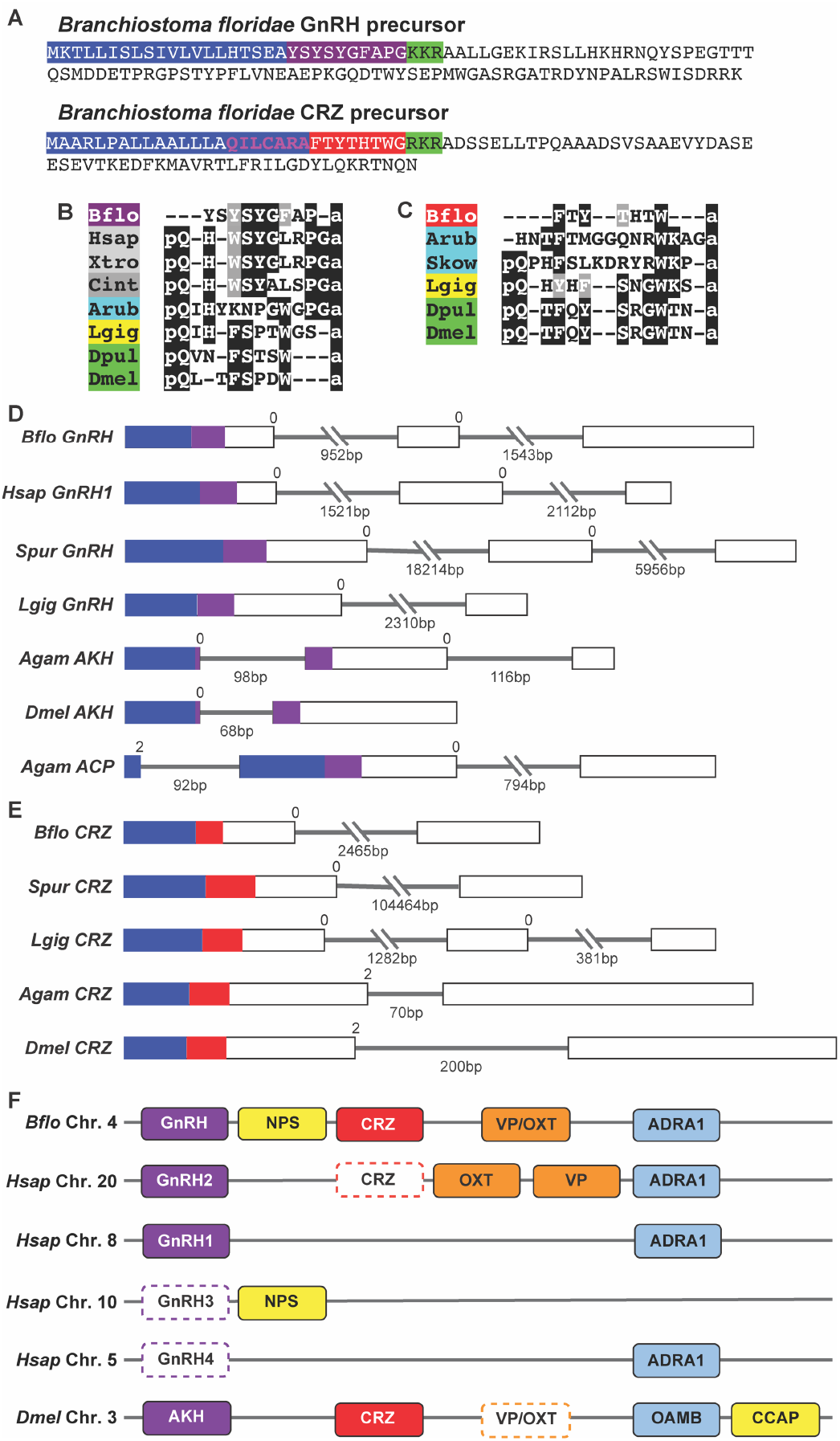
Identification of GnRH and CRZ precursors in the lancelet *Branchiostoma floridae*. **(A)** Amino acid sequences of *B. floridae* GnRH and CRZ neuropeptide precursors. Signal peptides are highlighted in blue, dibasic cleavage sites are highlighted in green and the putative mature peptides (without post-translational modifications) are highlighted in purple (GnRH) and red (CRZ). Sequence in pink within the CRZ signal peptide is part of the N’-terminally extended CRZ (CRZ-ext) reported by Roch et al. (2014). Multiple sequence alignments of **(B)** GnRH and AKH mature peptides, and **(C)** CRZ mature peptides. Comparisons of the exon-intron structure of genes encoding **(D)** GnRH, AKH and ACP precursors, and **(E)** CRZ precursors. Boxes represent exons (drawn to scale) and lines represent introns, with length underneath. Regions encoding signal peptides (blue) and mature peptides (purple and red) have been colored. The intron phase is indicated above the exonintron boundary. In deuterostomes, GnRH precursors are encoded by three exons and CRZ precursors are encoded by two exons, which distinguishes the two neuropeptides. **(F)** Conserved synteny for the genomic regions containing GnRH/AKH and CRZ precursor genes. Same colors are used for orthologs. For simplicity and ease of comparison, neighboring genes are aligned, locations of genes omitted and major gene loses indicated using broken boxes. Abbreviations: NPS, Neuropeptide S; CCAP, Crustacean cardioactive peptide; VP, Vasopressin; OXT, Oxytocin; ADRA1, Adrenoceptor Alpha 1; OAMB, Octopamine receptor in mushroom bodies. Species names: Agam, *Anopheles gambiae;* Arub, *Asterias rubens*; Bflo, *Branchiostoma floridae*; Cint, *Ciona intestinalis*; Dpul, *Daphnia pulex*; Dmel, *Drosophila melanogaster*; Hsap, *Homo sapiens*; Lgig, *Lottia gigantea*; Skow, *Saccoglossus kowalevskii*; Spur, *Strongylocentrotus purpuratus*; Xtro, *Xenopus tropicalis*.

Since both the putative *B. floridae* GnRH and CRZ mature peptides appear highly divergent, sequence alignments (Figure 1B and C) are not entirely suitable to infer homology. Therefore, we compared other sequence features, including the position of the mature peptide within the precursor and conserved introns, since these tend to be conserved in neuropeptide orthologs across diverse animal phyla (Mirabeau and Joly 2013). A comparison of GnRH (Figure 1D) and CRZ (Figure 1E) precursors from different animals shows that the mature peptide is flanked by a signal peptide at the N-terminus for both neuropeptide families. Moreover, the phase of the first intron is also 0 in both *GnRH* and *CRZ* genes from deuterostomes. However, a feature that distinguishes deuterostomian *GnRH* from deuterostomian *CRZ* sequences is the number of introns. *GnRH* from *B. floridae* and other deuterostomes have two introns interrupting the protein-coding sequence (Figure 1D) whereas deuterostomian *CRZ* sequences, including the one from *B. floridae*, have a single conserved intron (Figure 1E). Remarkably, synteny analysis indicates that the genomic regions containing GnRH/AKH and CRZ genes are largely conserved between *B. floridae* and *D. melanogaster* (Figure 1F). Taken together, sequence analyses indicates that the previously identified *B. floridae* precursor is the CRZ precursor. More importantly, our newly identified *B. floridae* precursor shares several features with GnRH/AKH precursors from other animals, and likely encodes the bona fide GnRH.

### Identification and functional characterization of *B. floridae* GnRH and CRZ receptors

Previously, two GnRH-like and two CRZ-like receptors were identified in the *B. floridae* genome (Mirabeau and Joly 2013; Roch, et al. 2014b). Several neuropeptide receptors have undergone masive expansions in *B. floridae* (On, et al. 2015; Wang, et al. 2017). Since a better, chromosome-level assembly for the *B. floridae* genome is now available (Simakov, et al. 2020), we questioned whether additional GnRH/CRZ-like receptors are present in *B. floridae*. To investigate this possibility, we performed a sensitive unbiased search for GnRH/CRZ-like receptors using clustering-based (Figure S3) and phylogenetic analyses. Our search indeed identified an additional receptor (Figures S4-S6) which grouped with the two previously identified *B. floridae* CRZ receptors (Figure 2A). Further, consistent with previous phylogenetic analyses (Roch, et al. 2014b; Tian, et al. 2016), the two GnRH-like receptors from *B. floridae* (Figure S7 and S8) grouped with GnRH/AKH/ACP-type receptors and the three *B. floridae* CRZ-like receptors grouped with other ambulacrarian and protostomian CRZ receptors. Hence, both the GnRH and CRZ receptor families in *B. floridae* have undergone independent expansions.

**Figure 2:**
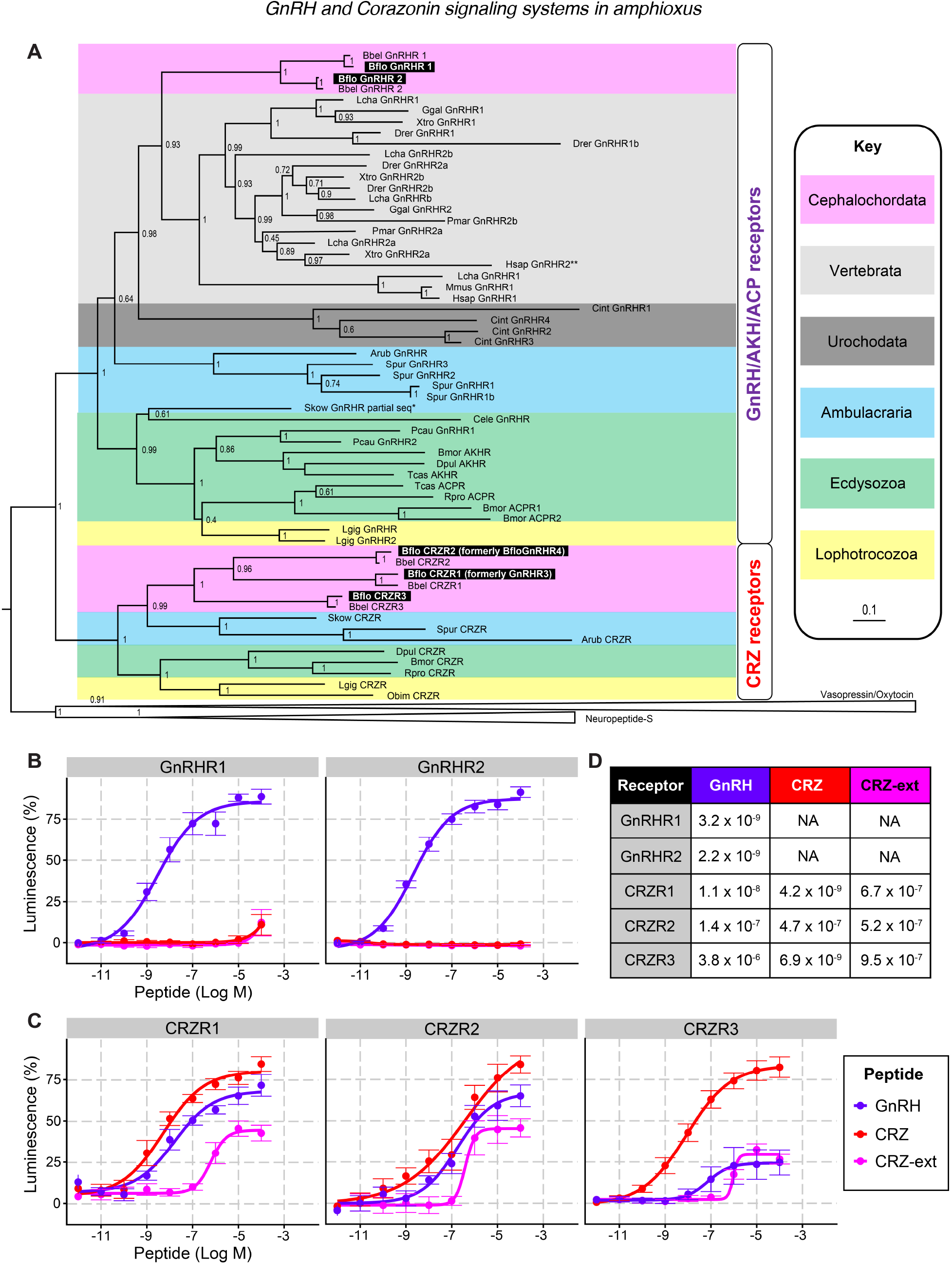
Phylogenetic analysis and functional characterization of GnRH and CRZ receptors in the lancelet *Branchiostoma floridae*. **(A)** Phylogenetic tree showing the relationship between GnRH and CRZ receptors. The tree was generated with PHYML 3.0 using the maximum-likelihood method. aLRT-SH-like support values (1000 replicates) are shown next to the nodes. The clade containing the Vasopressin/Oxytocin and Neuropeptide-S receptors was used to root the tree. Pastel-colored backgrounds represent different taxonomic groups (see key). Receptors functionally characterized in this study are highlighted in black. Species names: Bflo, *Branchiostoma floridae*; Bbel, *Branchiostoma belcheri*; Lcha, *Latimeria chalumnae*; Ggal, *Gallus gallus*; Xtro, *Xenopus tropicalis*; Drer, *Danio rerio*; Pmar, *Petromyzon marinus*; Hsap, *Homo sapiens*; Mmus, *Mus musculus*; Cint, *Ciona intestinalis*; Arub, *Asterias rubens*; Spur, *Strongylocentrotus purpuratus*; Skow, *Saccoglossus kowalevskii*; Cele, *Caenorhabditis elegans*; Pcau, *Priapulus caudatus*; Bmor, *Bombyx mori*; Dpul, *Daphnia pulex*; Tcas, *Tribolium castaneum*; Rpro, *Rhodnius prolixus*; Lgig, *Lottia gigantea*; Obim, *Octopus bimaculoides*. Functional characterization of *B. floridae* receptors belonging to the **(B)** GnRH/AKH/ACP receptor and **(C)** CRZ receptor clades in HEK293-G5a cells. GnRHR1 and GnRHR2 are only activated by GnRH. CRZR1, CRZR2 and CRZR3 are all activated by CRZ, GnRH and CRZ-ext; however, the receptors are most-sensitive and/or responsive to CRZ compared to GnRH and CRZ-ext. **(D)** EC_50_ values (M) for the dose-response curves in **B** and **C** illustrating the effectiveness of various ligands on *B. floridae* GnRH and CRZ receptors.

Having identified the *B. floridae* GnRH/CRZ-like peptides and receptors, we next sought to functionally validate these signaling systems by confirming that the putative GnRH and CRZ mature peptides can indeed activate their corresponding receptors. For this purpose, we expressed individual receptors in a heterologous expression system comprising Human embryo kidney 293 cells (HEK293-G5a) cells and monitored luminescence responses to three synthetic peptides. We tested the newly identified *B. floridae* GnRH (YSYSYGFAP-NH_2_), the CRZ (FTYTHTW-NH_2_), as well as the N-terminally-extended CRZ (CRZ-ext; pQILCARAFTYTHTW-NH_2_) predicted and tested previously (Roch, et al. 2014b). First, we tested the *B. floridae* GnRH receptors (GnRHR1 and GnRHR2) which were both specifically activated by GnRH in a dose-dependent manner, with EC_50_ values for the response in the low nanomolar range (Figure 2B and D). Neither of these two receptors were activated by CRZ or CRZ-ext. Together, these results confirm that the precursor identified here encodes the elusive *B. floridae* GnRH. Next, we tested the three CRZ receptors (CRZR1, CRZR2 and CRZR3). Consistent with Roch et al. (2014), CRZR2 (formerly GnRHR3) was activated by CRZ-ext, with a comparable EC_50_ value in the high nanomolar range (Figure 2C and D). Additionally, CRZR1 and CRZR3 were also activated by CRZ-ext; however, the EC_50_ values for the response were equally high. In contrast, the shorter CRZ activated all three CRZ receptors with a higher potency and efficacy compared to CRZ-ext (Figure 2C and D). Thus, CRZ rather than CRZ-ext, appears to be the endogenous ligand for the three *B. floridae* CRZ receptors. Identification of endogenous peptides in *B. floridae* via mass spectrometry can clarify whether CRZ, CRZ-ext or both are found *in vivo*. Surprisingly, GnRH was also able to activate all three CRZ receptors, with EC_50_ values for CRZR1 and CRZR2 responses in the mid to low nanomolar physiological range. Hence, there is a potential for cross-talk between the *B. floridae* GnRH and CRZ signaling systems *in vivo*. Decreased selection pressure on both the CRZ peptide and the ligand-binding pocket of CRZ receptors could explain how this signaling system was lost in vertebrates and other animals (Figure 3).

**Figure 3:**
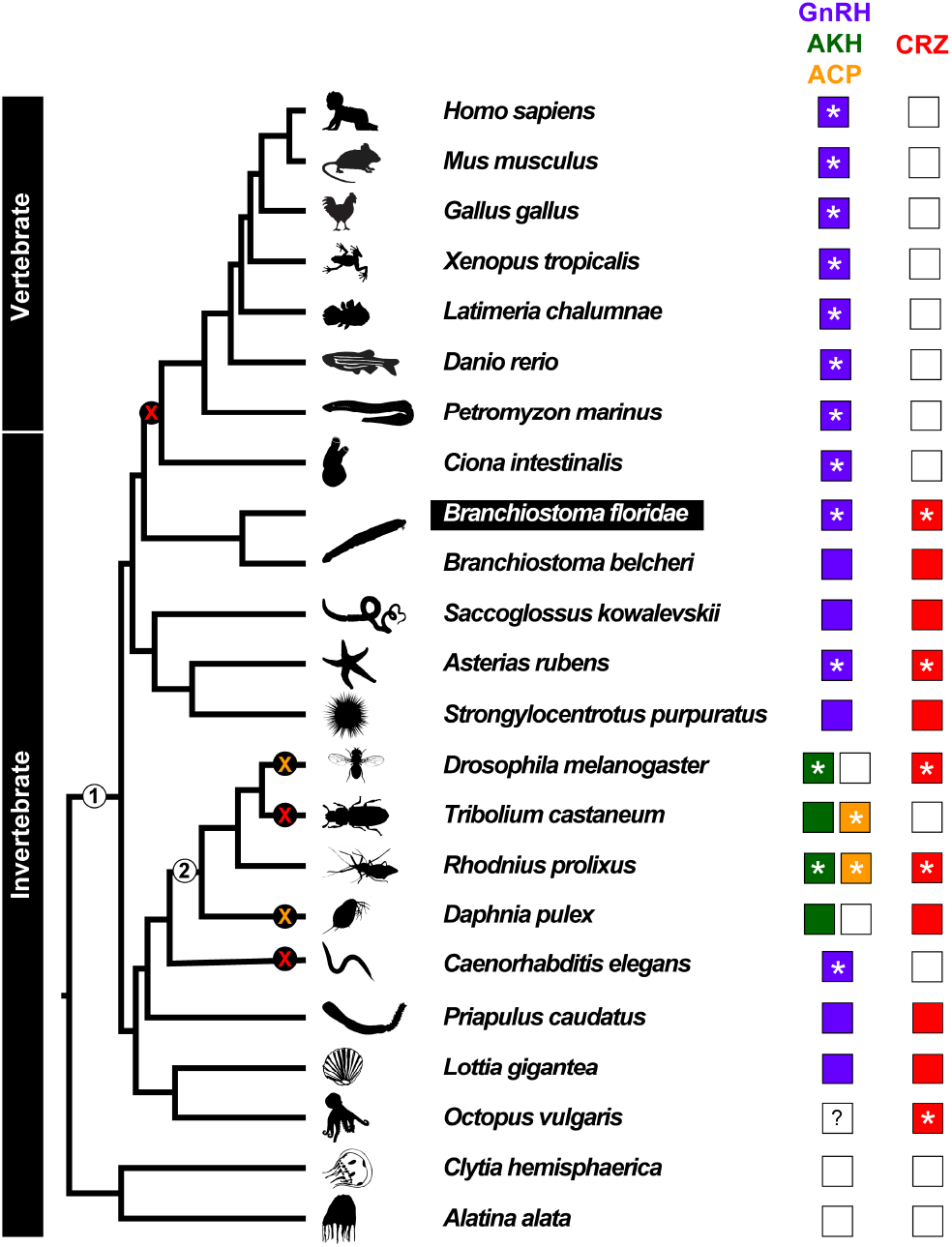
Evolution of GnRH and CRZ signaling systems. Animal phylogeny showing the occurrence of GnRH (purple), AKH (green), ACP (yellow) and CRZ (red) signaling systems. An empty box denotes absence of the signaling system. An asterisk indicates species in which the receptors have been functionally characterized and the question mark indicates uncertainty about the presence of that receptor in the genome. ACP (orange cross) and CRZ (red cross) systems have been independently lost in multiple lineages. Two major gene duplication events are marked in the phylogeny with numbers: (1) GnRH and CRZ signaling systems arose in a common ancestor of the Bilateria and (2) AKH and ACP signaling systems arose by duplication of the GnRH signaling system in a common ancestor of the Arthropoda. The lancelet *Branchiostoma floridae*, an invertebrate, is the closest relative of vertebrates to possess both the GnRH and CRZ signaling systems. Given the low quality of *Petromyzon marinus* genome assembly (Supplementary file 5), we cannot rule out the possibility that CRZ signaling is also found in vertebrates.

## Conclusion

In conclusion, we have discovered the bona fide GnRH, CRZ and their cognate receptors in the amphioxus, *B. floridae*. This suggests that CRZ signaling system has likely been lost in olfactores (vertebrates and urochordates). We acknowledge that the putative *B. floridae* GnRH and CRZ mature peptides require further validation using mass spectrometry. Discovery of both the signaling systems in one of the closest living relatives of vertebrates provides a framework to discover the physiological roles of GnRH and CRZ at the transition from invertebrates to vertebrates. Lastly, there is a strong need for neuropeptide discovery in diverse species as neuropeptide-encoding genes serve as excellent markers for single-cell transcriptome analyses (Ma, et al. 2021). Hence, our methodological approach highlighted here can facilitate these studies since it is ideal for identifying highly-divergent neuropeptides which evade typical BLAST searches.

## Data availability

The data underlying this article have been provided in the supplementary files. The scripts for dose-response curve analyses are available at github (https://github.com/Imnotabioinformatician/Branchiostoma_Dose_response_curves).

## Acknowledgements

The authors would like to thank Dr. Theresa McKim (University Hospital Würzburg) and Irina Wenzel (University of Würzburg) for helpful feedback during preparation of this paper. This work was supported by funding from the University of Würzburg (awarded to M.Z.) and the BBSRC Discovery fellowship (BB/W010305/1 awarded to L.A.Y.G.). This paper was typeset with the bioRxiv word template by @Chrelli: www.github.com/chrelli/bioRxiv-word-template.

## Author contributions

L.A.Y.G. and M.Z. conceived the study, performed computational analyses and analyzed the data. L.A.Y.G. performed the experimental work. M.Z. and L.A.Y.G. wrote the manuscript.

## Materials and Methods

### Identification of the *B. floridae* GnRH precursor using a Hidden Markov Model search

A GnRH-like precursor from *B. floridae* was identified using a custom HMM for GnRH/CRZ precursors. The sequences used to generate the HMM (Supplementary file 1) were obtained from publicly-available databases (NCBI). These sequences were aligned using the automated option in MUSCLE (Edgar 2004) and the resulting alignment trimmed using TrimAL (Capella-Gutierrez, et al. 2009). This trimmed alignment was used to generate the HMM (Supplementary file 2) using HMMER3.0 (http://hmmer.org/) (Eddy 2011). This model was used to search *Branchiostoma floridae* and *Branchiostoma belcheri* proteomes with an e-value of 1e-1.

### Sequence analyses and alignments

Potential signal peptide cleavage sites in *B. floridae* GnRH and CRZ precursors were predicted using SignalP 6 Server (Teufel, et al. 2022). GnRH and CRZ mature peptide sequences were aligned using Clustal Omega (Sievers, et al. 2011). The alignments were adjusted and shaded manually. Gene structures of GnRH and CRZ precursors were predicted using Webscipio 2.0 (Hatje, et al. 2011). Membrane topology of the receptors was predicted *in silico* using Protter and DeepTMHMM (https://dtu.biolib.com/DeepTMHMM) (Omasits, et al. 2014). Synteny analysis of *GnRH/AKH* and *CRZ* was based on the one performed earlier by Roch et al., 2014 (Roch, et al. 2014a). *Drosophila* gene loci were obtained from FlyBase (Gramates, et al. 2022) and *B. floridae CRZ* loci was determined via genome BLAST.

### Clustering and phylogenetic analyses of GnRH and CRZ receptors in Bilateria

Putative *B. floridae* GnRH and CRZ receptors were identified using a combination of clustering-based and phylogenetic analyses. Briefly, GnRH and CRZ receptor sequences (Supplementary file 3) were aligned using MUSCLE and trimmed using TrimAL as described above. This alignment was used to generate a custom HMM for GnRH and CRZ receptors (Supplementary file 4) with HMMER3.0. We used this model to search for GnRH/CRZ receptors (with an e-value of 1e-25) from select species belonging to major animal phyla. Predicted proteomes of the following species were searched: *Asterias rubens, Bombyx mori, Caenorhabditis elegans, Ciona intestinalis, Clytia hemisphaerica, Danio rerio, Daphnia pulex, Gallus gallus, Homo sapiens, Latimeria chalumnae, Lottia gigantea, Mus musculus, Octopus bimaculoides, Petromyzon marinus, Priapulus caudatus, Rhodnius prolixus, Saccoglossus kowalevskii, Strongylocentrotus purpuratus, Tribolium castaneum* and *Xenopus tropicalis*. Since predicted proteomes of *Alatina alata, Branchiostoma belcheri* and *Branchiostoma floridae* were not available, their transcriptomes were first translated into protein sequences (minimum length of 100 amino acids) using the prediction option in TransDecoder (http://transdecoder.github.io/) before performing the search. The sources for these predicted proteomes and transcriptomes are available in Supplementary file 5. Independently, the GnRH and CRZ receptor sequences used for generating the custom HMM above were also used as a query to perform a BLASTp search with an e-value cutoff of 1e-15. The resulting hits from the HMM and BLASTp searches were merged. Redundant sequences were then eliminated (at a 98% threshold) using CD-Hit (Fu, et al. 2012). These searches retrieved several receptors including those activated by other neuropeptides and monoamines. To identify putative GnRH and CRZ receptors, all the receptor sequences were clustered in CLANS (https://toolkit.tuebingen.mpg.de/tools/clans) (Frickey and Lupas 2004), using the BLOSUM 62 matrix and extracted BLAST High scoring pairs with an e-value of 1e-15 as a threshold. The CLANS analysis file is available in Supplementary file 6. To identify the different clusters, we used the linkage-clustering option with at least 2 links. Receptors sequences from GnRH/CRZ, Vasopressin/Oxytocin and Neuropeptide-S/NGFFFamide clusters were extracted (Supplementary file 7) and used for phylogenetic analysis. Sequences were aligned using the iterative refinement method E-INS-i in MAFFT version 7.0 (Katoh, et al. 2019). The alignment was trimmed with TrimAl in gappy mode (Capella-Gutiérrez, et al. 2009). The maximum-likelihood tree was produced using PHYML (ngphylogeny.fr) with the LG general amino-acid replacement matrix and 4 discrete gamma models (Le and Gascuel 2008). Branch support was calculated based on 1000 replicates with the aLRT-SH-like methodology (Guindon, et al. 2010). The trimmed sequences used for the phylogenetic analysis are available in Supplementary file 8.

### Functional characterization of *B. floridae* GnRH and CRZ receptors

*Branchiostoma floridae* GnRH (YSYSYGFAP-NH2), CRZ (FTYTHTW-NH2) and CRZ-ext (pQILCARAFTYTHTW-NH2) were custom synthesized by NovoPro Bioscience Inc with a purity of >95%. 5 mM stock solutions of all peptides were first prepared in 50% DMSO. Two putative GnRH receptors and three putative CRZ receptors were identified in *B. floridae* based on our clustering and phylogenetic analyses. These receptors were codon-optimized for mammalian cell lines, synthesized and cloned into pcDNA3.4(+) vector by Genscript synthesis services. A 5’ partial Kozak translation initiation sequence (CCACC) was also included to improve expression. The codon-optimized receptor sequences are provided in the Supplementary file 9. HEK293-G5a (Angio-proteomie CAT no. cAP0200GFP-AEQ-Cyto) cells were cultured in 96 well-plates containing 100μl of DMEM (Thermo; Cat. No. 10566016) supplemented with 10% foetal bovine serum (Thermo; Cat. No. 10082147). The plates were kept in an incubator at 37 °C and 5 % CO_2_. Upon reaching 85% confluency, cells were co-transfected with plasmids encoding individual receptors or the empty pcDNA3.1+ vector (control), and the Gαqi9 promiscuous protein [Addgene; Cat. No. 125711 (Masharina, et al. 2012)], as described previously (Yanez-Guerra, et al. 2022). Transfections were carried out with 60 ng of each plasmid and 0.45 μl of the transfection reagent Transfectamine 5000 (AAT-bioquest; Cat. No. 60022) per well. Two days post-transfection, the media was replaced with fresh DMEM medium supplemented with 4 mM coelenterazine-H (Thermo Fisher Scientific; Cat. No. C6780). The cells were incubated for 2 hrs and then used for the luminescence assay. Various concentrations of the three synthetic peptides ranging from 10^−4^ M to 10^−12^ M were prepared in DMEM-media and tested on the cells. Luminescence levels were recorded and integrated over a 65-second measurement period in a FlexStation 3 Multi-Mode Microplate Reader (Molecular Devices). A minimum of two independent transfections (biological replicates) for each receptor and two-three technical replicates for each peptide concentration were performed. The responses were normalized to the maximum response obtained following the addition of peptide in each experiment (100% activation) and to the response obtained with the vehicle media (0% activation). Normalised data from the independent transfections was used to plot the dose-response curves (fitted with a four-parameter curve) using the package drc in R (Ritz, et al. 2015). The raw data for the emtpy-vector control and receptor characterization are provided in Supplementary files 10 and 11, respectively. The script used for the dose-response curve analysis is available on Github (https://github.com/Imnotabioinformatician/Branchiostoma_Dose_response_curves).

**Figure S1:**
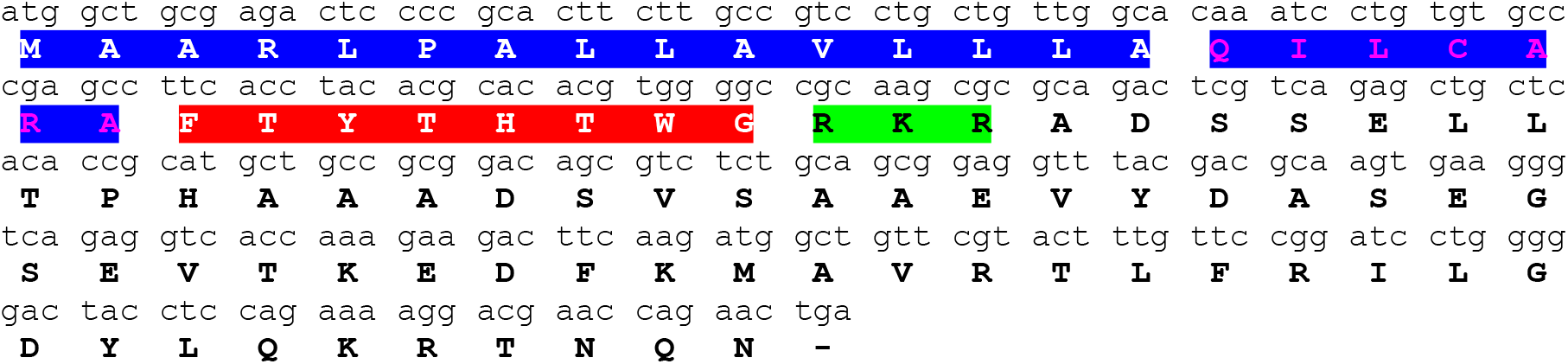
*Branchiostoma floridae* CRZ cDNA sequence (Accession no: KF601546.1) and the deduced amino acid sequence. Within the amino acid sequence, the signal peptide is highlighted in blue, the mature peptide is in red and the predicted cleavage site is in green. The residues in pink within the signal peptide were part of the putative CRZ mature peptide predicted previously by Roch et al. (2014).

**Figure S2:**
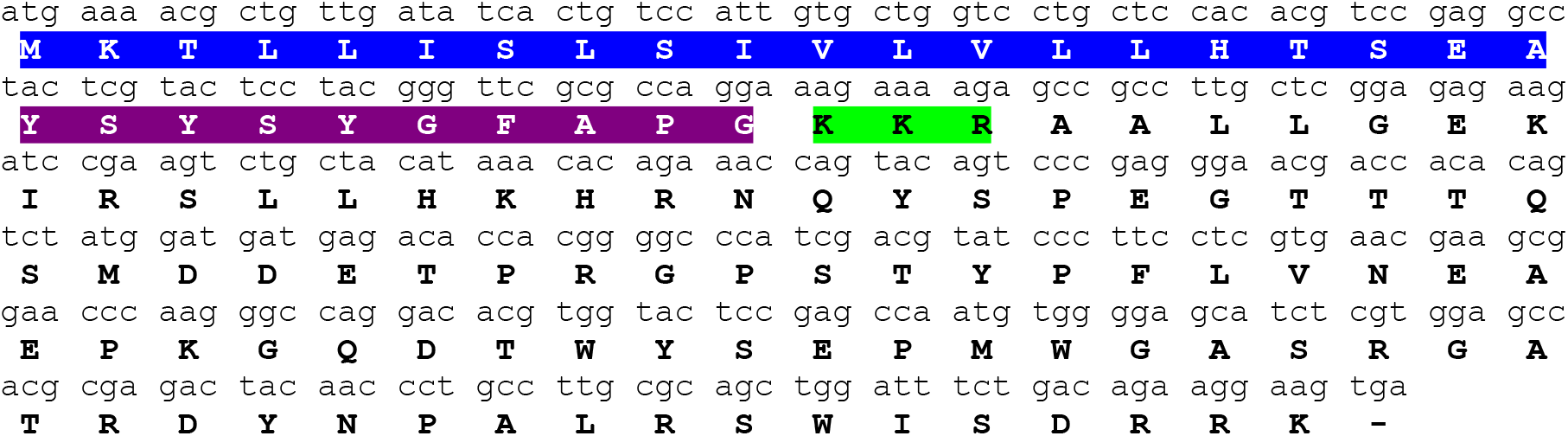
*Branchiostoma floridae* GnRH cDNA sequence (Accession no: XM_035819417.1) and the deduced amino acid sequence. Within the amino acid sequence, the signal peptide is highlighted in blue, the mature peptide is in purple and the predicted cleavage site is in green.

**Figure S3:**
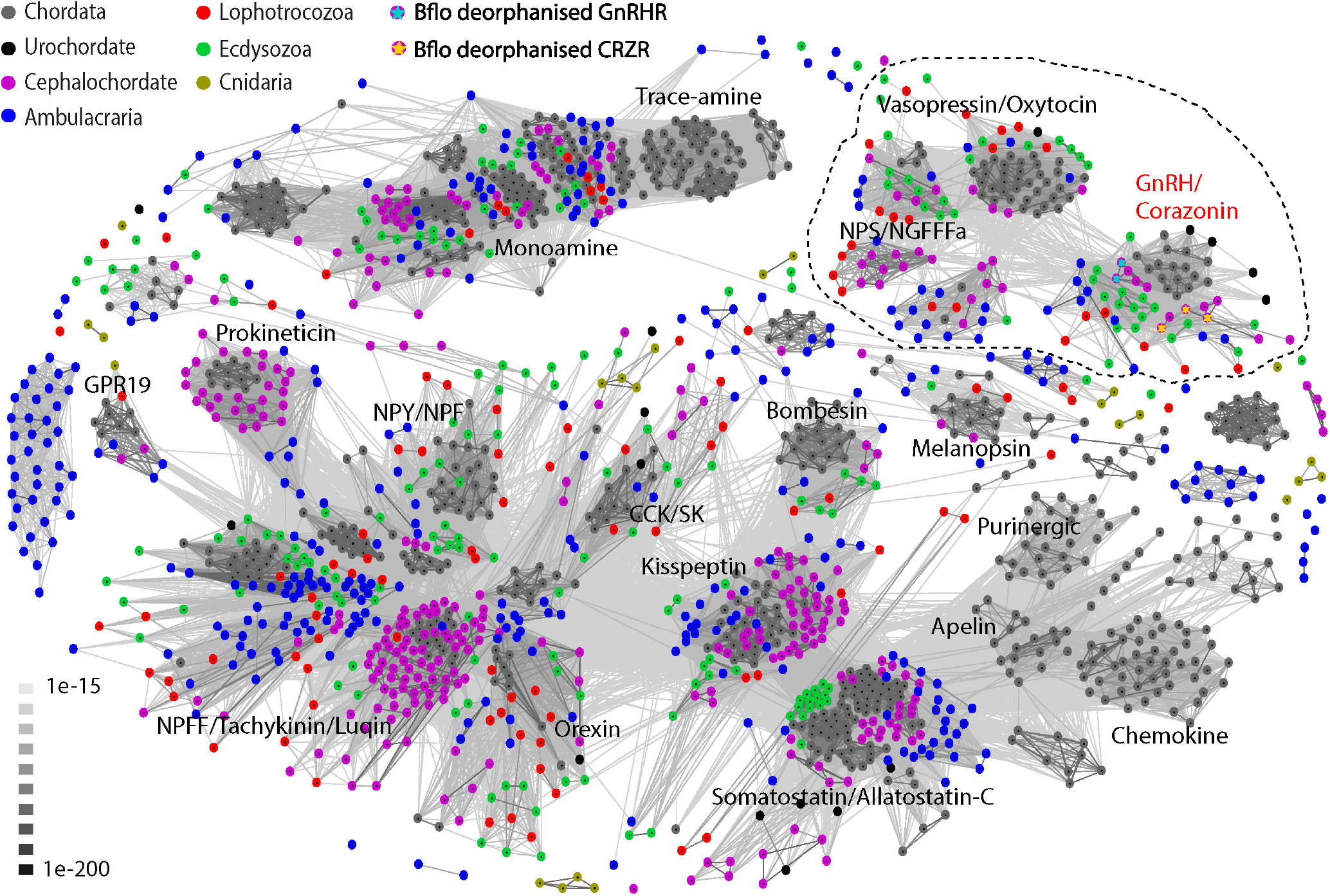
Cluster analysis of GPCRs shows that GnRH and CRZ receptors cluster closely with vasopressin/oxytocin and Neuropeptide-S (NPS)/NGFFFa receptors (dotted region). Each dot represents an individual receptor which has been color-coded according to its phyla. The receptors in the dotted region were used for the phylogenetic analysis in Figure 2A. The three CRZ receptors and two GnRH receptors from *Branchiostoma floridae* characterized in this study have been marked with yellow and cyan stars, respectively. Edges represent BLAST connections of *P* value < 1e-15.

**Figure S4:**
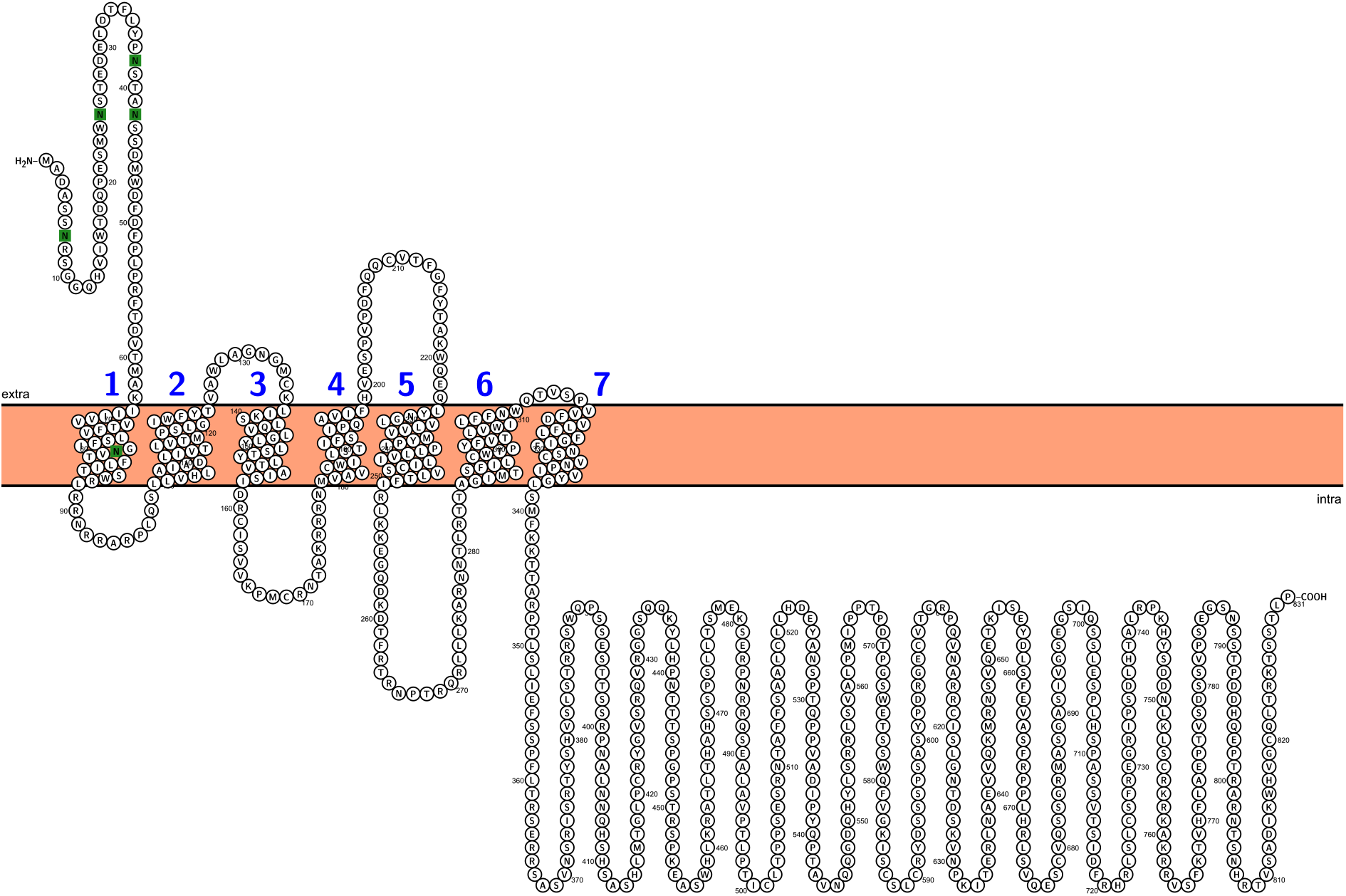
*In silico* prediction of the *Branchiostoma floridae* CRZR1 topology. The transmembrane domains are numbered successively in blue and the putative N-glycosylation sites are shown with green boxes.

**Figure S5:**
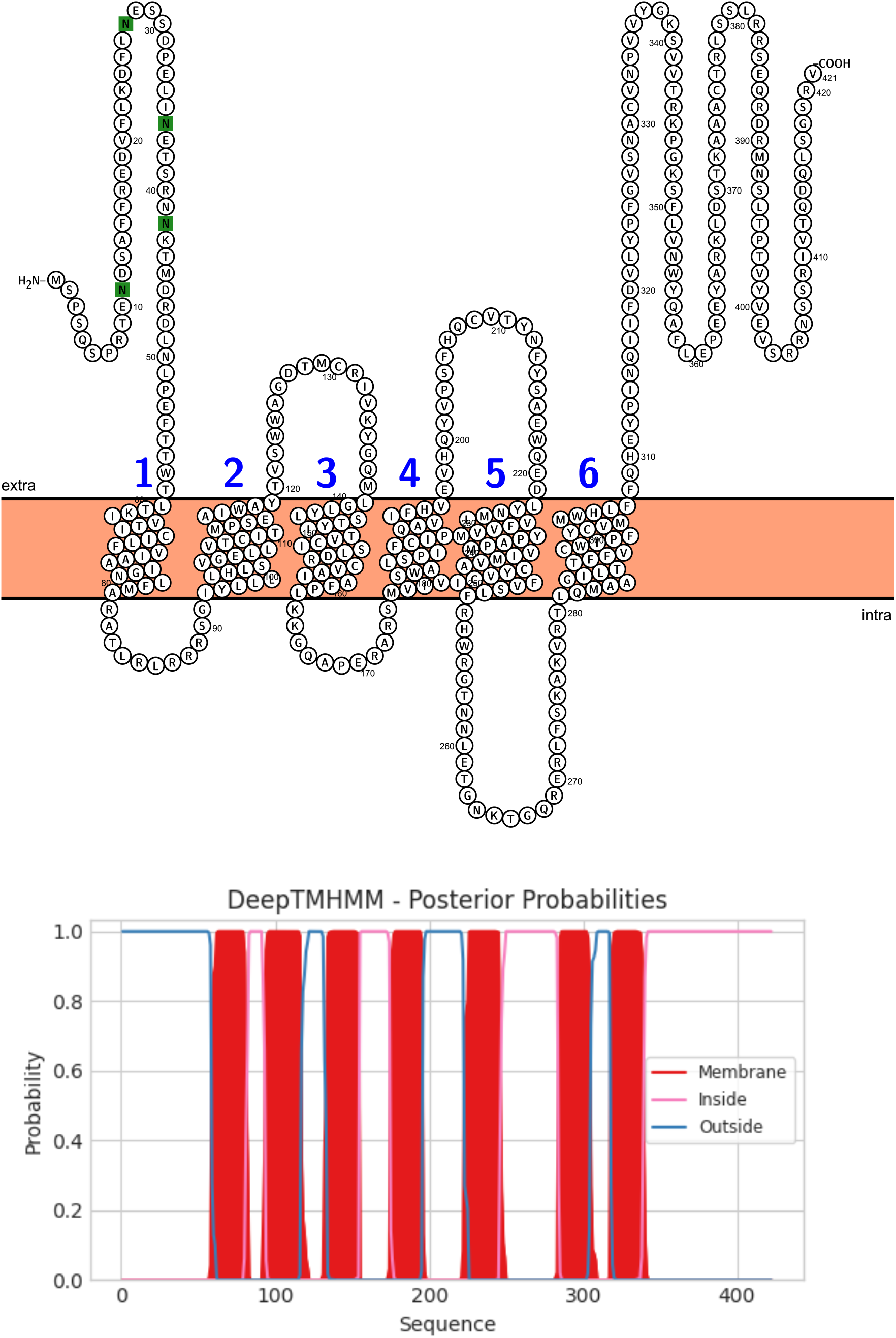
*In silico* prediction of the *Branchiostoma floridae* CRZR2 topology using Protter (top) and DeepTMHMM (bottom). For the prediction using Protter, the transmembrane domains are numbered successively in blue and the putative N-glycosylation sites are shown with green boxes. Since GPCRs typically have 7 transmembrane domains, the topology predicted by DeepTMHMM appears to be more accurate than Protter for this receptor.

**Figure S6:**
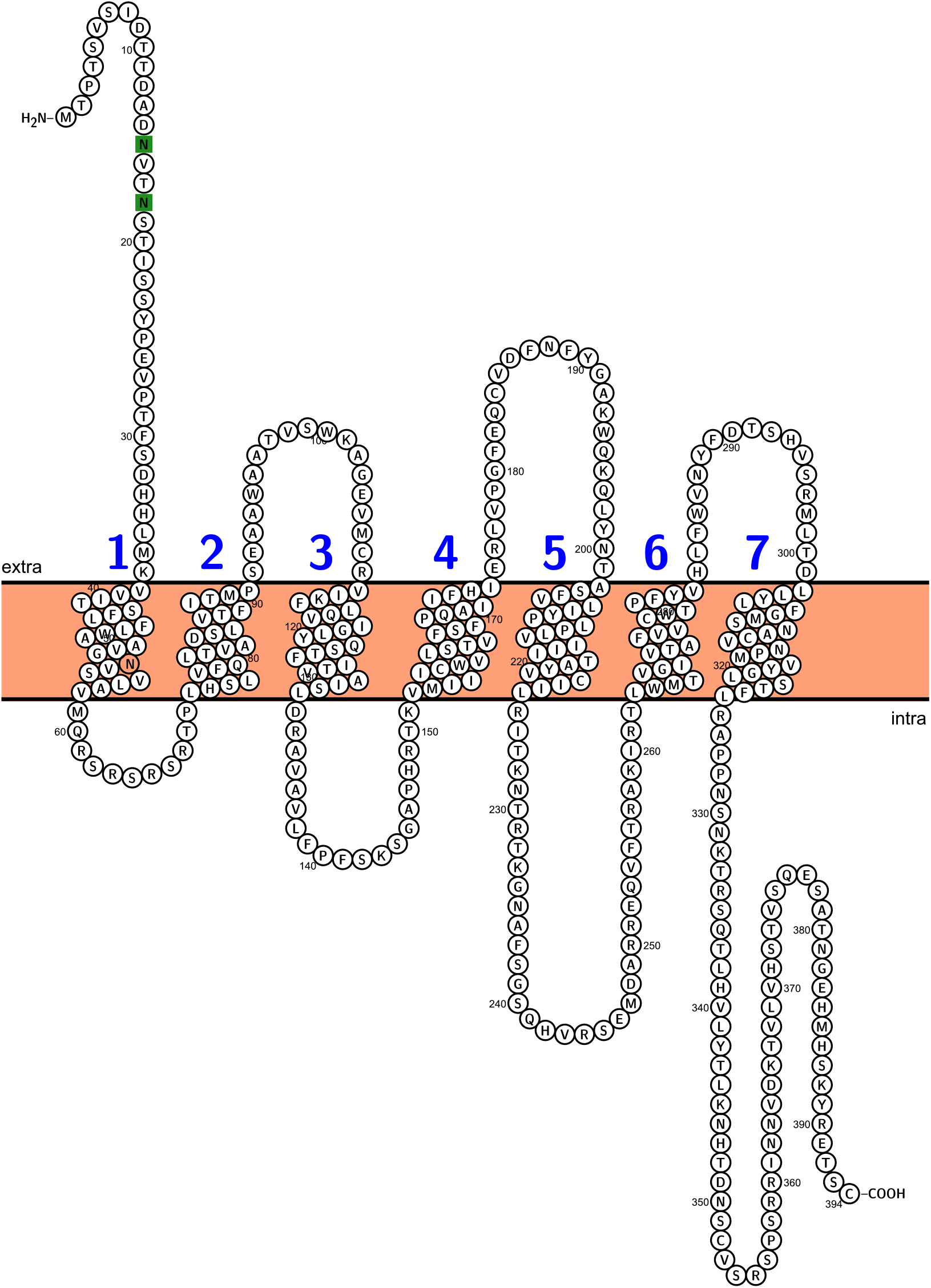
*In silico* prediction of the *Branchiostoma floridae* CRZR3 topology. The transmembrane domains are numbered successively in blue and the putative N-glycosylation sites are shown with green boxes.

**Figure S7:**
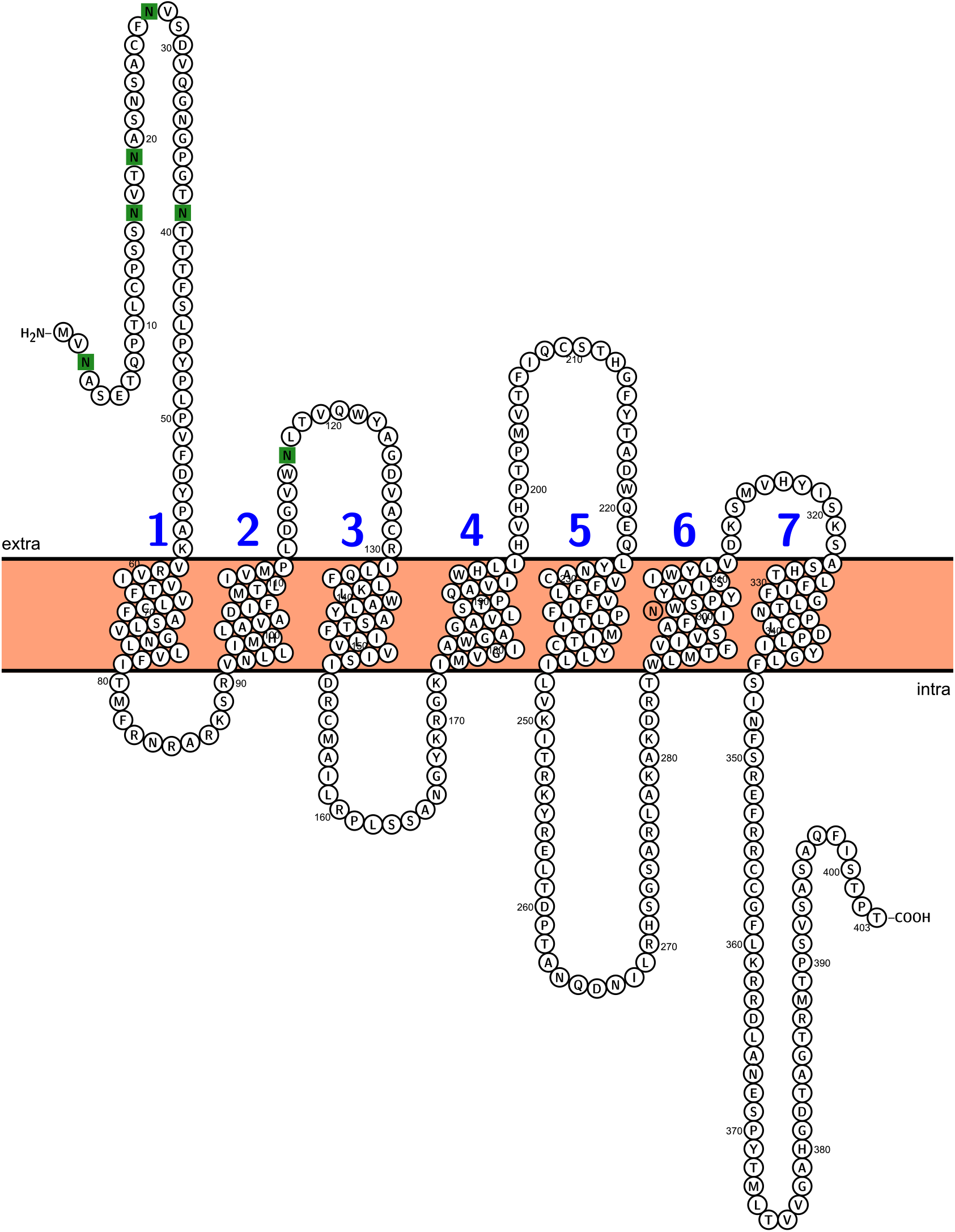
*In silico* prediction of the *Branchiostoma floridae* GnRHR1 topology. The transmembrane domains are numbered successively in blue and the putative N-glycosylation sites are shown with green boxes.

**Figure S8:**
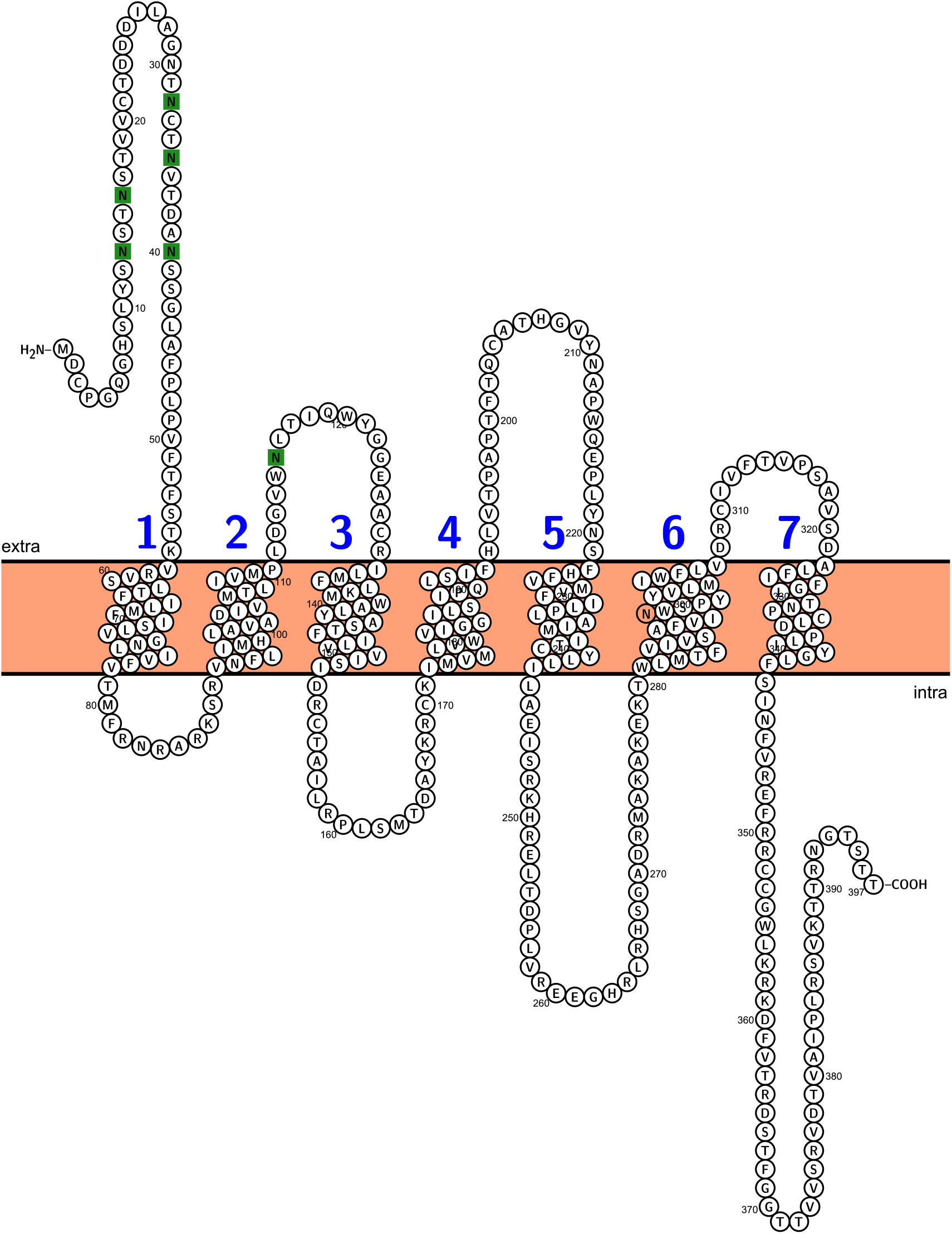
*In silico* prediction of the *Branchiostoma floridae* GnRHR2 topology. The transmembrane domains are numbered successively in blue and the putative N-glycosylation sites are shown with green boxes.

