## Supplementary material for "Discovery of paralogous GnRH and corazonin signaling systems in an invertebrate chordate": SupplementaryFile9_codon_optimized_Branchiostoma_receptors.docx

**Supplementary File 9**

Codon optimized sequence of the *Branchiostoma floridae* receptors GNRHR1, GNRHR2, CRZR1, CRZR2 and CRZR3. The region highlighted corresponds to an added Kozak sequence to improve the expression.

>Branchiostoma_floridae_GnRHR1_ACC68665_1_XP_035657822.1__EU433377.1

CCACCATGGTGAATGCCAGCGAGACACAGCCAACCCTGTGCCCCAGCTCCAACGTGACCAATGCCTCCAACTCTGCCTGTTTTAACGTGTCTGACGTGCAGGGCAATGGCCCAGGCACAAACACCACAACCTTTAGCCTGCCCTACCCTCTGCCAGTGTTCGATTATCCCGCCAAGGTGAGAGTGATCGTGACCTTCGTGCTGTGCTTTGCCTCCCTGGTGGGCAATCTGCTGGTGTTCATCACAATGTTTCGGAATAGAGCCAGGAAGTCTAGGGTGAACCTGCTGATCATGCACCTGGCCGTGGCCGACATCTTCATGACCCTGATCGTGATGCCTCTGGATGGCGTGTGGAACCTGACAGTGCAGTGGTACGCAGGCGACGTGGCATGCAGAATCCTGCAGTTTCTGAAGCTGTGGGCCCTGTATGCCAGCACCTTCATCCTGGTGGTCATCTCCATCGATAGATGTATGGCCATCCTGAGGCCCCTGTCTAGCGCCAATGGCTACAAGAGGGGCAAGATCATGGTGGGAATCGCATGGGGAGCAGGAGCCGTGCTGTCTACCCCTCAGGCCGTGATCTGGCACCTGATCCACGTGCACCCCACCCCTATGGTGACCTTCATCCAGTGCTCCACCCACGGCTTCTACACAGCCGACTGGCAGGAGCAGCTGTATAACGCCTGCGTGTTCTTTCTGGTGTTCATCTTTCCTCTGACCATCATGATCACATGTTACCTGCTGATCCTGGTGAAGATCACCCGCAAGTATCGGGAGCTGACAGACCCAACCGCCAATCAGGATAACATCCTGAGGCACAGCGGCTCCGCCAGGCTGGCCAAGGCCAAGGATAGAACCTGGCTGATGACATTTGTGATCGTGAGCGCCTTCGTGATCAATTGGTCCCCATACTATGTGATCTCTATCTGGTACCTGGTGGACAAGTCCATGGTGCACTATATCTCTAAGAGCGCCTCCCACACCCTGTTCATCTTTGGCCTGACAAATCCTTGTCTGGACCCCCTGATCTACGGCCTGTTCTCTATCAACTTTAGCCGCGAGTTCCGGAGATGCTGTGGCTTTCTGAAGAGGCGCGACCTGGCCAACGAGAGCCCCTATACAATGCTGACCGTGGTGGGAGCACACGGCGATACAGCAGGAACCCGGATGACACCCTCTGTGAGCGCCTCCGCCCAGTTCATCTCCACACCTACCTGA

>Protein_Branchiostoma_floridae_GnRHR1_ACC68665_1_XP_035657822.1__EU433377.1

MVNASETQPTLCPSSNVTNASNSACFNVSDVQGNGPGTNTTTFSLPYPLPVFDYPAKVRVIVTFVLCFASLVGNLLVFITMFRNRARKSRVNLLIMHLAVADIFMTLIVMPLDGVWNLTVQWYAGDVACRILQFLKLWALYASTFILVVISIDRCMAILRPLSSANGYKRGKIMVGIAWGAGAVLSTPQAVIWHLIHVHPTPMVTFIQCSTHGFYTADWQEQLYNACVFFLVFIFPLTIMITCYLLILVKITRKYRELTDPTANQDNILRHSGSARLAKAKDRTWLMTFVIVSAFVINWSPYYVISIWYLVDKSMVHYISKSASHTLFIFGLTNPCLDPLIYGLFSINFSREFRRCCGFLKRRDLANESPYTMLTVVGAHGDTAGTRMTPSVSASAQFISTPT

>Branchiostoma_floridae_GnRHR_2_ACC68666_1_XP_035668163.1_EU433378.1

CCACCATGGATTGCCCTGGCCAAGGACACAGTCTTTACTCAAACTCTACTAACTCCACAGTAGTGTGTACAGACGATGACATACTGGCGGGAAATACTAATTGTACCAACGTCACCGATGCCAATTCGTCAGGCTTGGCCTTTCCCTTACCAGTCTTCACTTTCTCAACGAAAGTCCGCGTCTCTTTGACATTTATCTTAATGTTCATATCTCTGGTGGGAAATCTCATCGTCTTCGTCACGATGTTCCGAAATCGCGCCAGGAAGTCTCGCGTGAACTTCCTCATCATGCACCTGGCCGTGGCCGATATCGTCATGACGTTGATCGTGATGCCGCTAGATGGCGTGTGGAACTTGACGATCCAGTGGTACGGTGGGGAGGCTGCGTGCCGCATTCTGATGTTCCTGAAGATGTGGGCGCTGTACGCGTCTACCTTTATTCTCGTGGTCATCAGTATTGACCGATGCACGGCCATCCTCCGTCCTCTGAGCATGACGGACGCCTACAAGAGATGCAAGATCATGGTCATGCTGGTGTGGGTCATCGGGGGCATCCTCAGCATCCCTCAGCTCTCCATTTTCCATCTCGTGACGCCAGCGCCGACCTTCACGCAGTGTGCGACACACGGAGTCTACAACGCCCCCTGGCAGGAGCCGTTGTACAACAGCTTCCACTTCGTCATGGTTTTCATCCTGCCGCTGGCCATCATGATCACGTGTTACCTCCTCATCCTGGCCGAAATCTCCCGCAAGCACCGGGAGCTGACTGATCCACTTGTCCGGGAGGAGGGGCACAGACTCCGCCACTCCGGTGCAGACCGCATGGCCAAGGCCAAGGAGAAGACCTGGCTGATGACCTTTGTCATCGTGTCGGCCTTCGTCATCAACTGGTCCCCGTACTACGTCCTGATGATCTGGTTCCTGGTGGATCGGTGCATCGTCTTCACCGTGCCGTCAGCGGTGAGCGACGCCCTCTTCATCTTCGGCCTGACCAACCCCTGCCTCGACCCGCTCATCTACGGCCTGTTCTCCATCAACTTCGTGCGGGAGTTCCGTCGCTGCTGCGGCTGGCTCAAGCGCAAGGACTTCGTGACCAGAGACAGCACCTTCGGCGGGACCACGGTCGTCAGCCGGGTCGACACCGTCGCCATCCCGCTACGATCGGTGAAGACCACTCGCAACGGAACATCAACCACCTGA

>Protein_Branchiostoma_floridae_GnRHR_2_ACC68666_1_XP_035668163.1_EU433378.1

MDCPGQGHSLYSNSTNSTVVCTDDDILAGNTNCTNVTDANSSGLAFPLPVFTFSTKVRVSLTFILMFISLVGNLIVFVTMFRNRARKSRVNFLIMHLAVADIVMTLIVMPLDGVWNLTIQWYGGEAACRILMFLKMWALYASTFILVVISIDRCTAILRPLSMTDAYKRCKIMVMLVWVIGGILSIPQLSIFHLVTPAPTFTQCATHGVYNAPWQEPLYNSFHFVMVFILPLAIMITCYLLILAEISRKHRELTDPLVREEGHRLRHSGADRMAKAKEKTWLMTFVIVSAFVINWSPYYVLMIWFLVDRCIVFTVPSAVSDALFIFGLTNPCLDPLIYGLFSINFVREFRRCCGWLKRKDFVTRDSTFGGTTVVSRVDTVAIPLRSVKTTRNGTSTT

>Branchiostoma_floridae_CRZR1_ACC68668_1_XP_035668126.1_XM_035812233.1 __(Formerly_GnRHR3)

CCACCATGGCCGATGCCAGCTCCAACAGAAGCGGCGGCCAGCACGTGATCTGGACAGACCAGCCTGAGAGCATGTGGAATTCCACAGAGGACGAGCTGGATACCTTCCTGTATCCAAACTCCACCGCCAATTCTAGCGACATGTGGGACTTCGATTTTCCACTGCCCAGGTTTACAGATGTGACCATGGCCAAGATCATCATCGTGGTGGTGACCTTCGTGCTGTCTTTTATCGGCAACGTGACCTTCCTGATCACAAGCTGGAGACTGCGGAGAAATAGGAGGGCAAGGCCACTGCAGAGCCTGCTGGTGCACCTGGCCATCGCCGATCTGATCGTGACACTGGTGACCATGCCCTCCCTGGGCATCTGGTTTTACACCGTGGCATGGCTGGCAGGAAACGGAATGTGCAAGCTGATCAAGTCCCTGCAGGTGCTGGGCCTGTACCTGTCTACATATCTGACCGTGGCCATCTCCATCGATCGCTGCATCTCTGTGGTGAAGCCTATGTGCCGGAACACAGCCAAGCGGAGAAGGAATATGGTGGCCGTGTGCTGGATTCTGTCTACCATCTTCAGCATCCCACAGGCCGTGATCTTTCACGTGGAGAGCCCTGTGCCAGACTTCCAGCAGTGCGTGACCTTCGGCTTCTACACAGCCAAGTGGCAGGAGCAGCTGTATAACGGCCTGGTGCTGGTGGTCATGTACCCCATCCCTCTGCTGGTCATCCTGATCTGTTCCGTGCTGACATTCATCAGACTGAAGAAGGAGGGCCAGGACAAGGATACATTTCGCACCCGGAATCCCACCAGACAGAGGCTGCTGCTGAAGGCCCGCAACAATACACTGCGGACCACAGCCGGCATCATGACCAGCTTCATCCTGTGCTGGACCCCCTATTTTGTGACCCTGGTGTGGATTCTGTTCTTCAACTGGCAGACCGTGTCCCCTGTGGTGTTCGACGTGCTGTTCCTGTTTGGCATCTTTAACAGCTGCGTGAATCCAATCGTGTACGGCCTGTCCATGTTCAAGAAGACCACAGCCAGACCAACACTGTCCCTGATCGAGTTCTCCTCTCCCTTTCTGACCAGGTCTGAGCGCCGGTCTGCCAGCGTGAACAGCAGAATCTCCAGGACATATAGCCACGTGTCCCTGTCTACCAGAAGGTCCTGGCAGCCTAGCTCCGAGTCTACCACATCTAGCAGACCAAATGCCCTGAACAATCAGCACAGCCACTCCGCCTCTCACCTGATGACCGGCCTGCCATGCAGATATGGCGTGAGCCGCCAGGTGCGGGGAGGCTCCCAGCAGAAGTACCTGCACCCAAACACCACAACCAGCCCAGGACCATCTACACGCAGCCCAAAGGAGGCCTCCTGGCACCTGAAGAGGGCCACACTGACCCACGCACACTCCTCTCCCTCCCTGCTGACCTCTATGGAGAAGAGCGAGAGGCCTAATCGCCGGCAGAGCGAGGCCCTGGCAGTGCCTACACTGCCAACCATCTGCCTGACACCCCCTAGCGAGTCCAGGAACACCGCCTTCTTCAGCGCCGCCCTGTGCCTGCTGCACGACGAGTACGCCAATAGCCCCACACAGCCACCCGTGGCCGATATCCCCTATCAGCCTACCGCCGTGAACCAGCAGGGCGACCAGCACTACCTGTCCAGAAGGCTGTCTGTGGCCCTGCCTATGATCCCTCCAACACCAGACACCCCAGGCAGCTGGGAGACAAGCTCCTGGCAGTTCGTGGGCAAGATCTCTTGCAGCCTGTGCAGATATGACTCTAGCTCCCCATCCGCCTCTTACCCTGATAGGGGAGAGTGCGTGACCGGCCGGCCCCAAGTGAATGCACGCCGGTGTATCAGCCTGGGCAACACAGATTCCAAAGTGAATCCTAAGATCACCGAGAGACTGAACGCCGAGGTGGTGCAGAAGATGAGGAATTCCGTGCAGGAGACCAAGATCTCCGAGTATGACCTGTCTTTCGAGGTGGCCAGCTTTAGACCCCCTCTGCACAGGCTGAGCGTGCAGGAGTCCTGCGTGCAGTCTAGCGGAAGGATGGCATCCGGAGCATCTATCGTGGGCAGCGAGGGCTCCATCCAGTCCTCTCTGGAGTCTCCTCTGCACAGCCCAGCCAGCTCCGTGACAAGCATCGATTTCCGCCACCGGCTGTCTCTGTGCAGCTTTAGAGAGGGCAGAATCCCCTCCGACCTGCACACCGCCCTGAGGCCTAAGCACTACTCTGACGATAACCTGAAGCTGAGCTGTAGAAAGAGGAAGGCCAAGAGAAGGGTGTCCTTCAAGACAGTGCACTTTCTGGCCGAACCTACCGTGTCTGATTCTAGCGTGCCATCCGAGGGCTCTAATTCCTCTACCCCAGACGATCACCAGGAGCCAACCCGCGCCAGGAACACATCTAATCACCGCACCGTGAGCGCCGACATCAAGTGGCACGTGGGCTGTCAGCTGACACGGAAGACCAGCTCCACACTGCCCTGAGAG

>Protein_Branchiostoma_floridae_CRZR1_ACC68668_1_XP_035668126.1_XM_035812233.1 __(Formerly_GnRHR3)

MADASSNRSGGQHVIWTDQPESMWNSTEDELDTFLYPNSTANSSDMWDFDFPLPRFTDVTMAKIIIVVVTFVLSFIGNVTFLITSWRLRRNRRARPLQSLLVHLAIADLIVTLVTMPSLGIWFYTVAWLAGNGMCKLIKSLQVLGLYLSTYLTVAISIDRCISVVKPMCRNTAKRRRNMVAVCWILSTIFSIPQAVIFHVESPVPDFQQCVTFGFYTAKWQEQLYNGLVLVVMYPIPLLVILICSVLTFIRLKKEGQDKDTFRTRNPTRQRLLLKARNNTLRTTAGIMTSFILCWTPYFVTLVWILFFNWQTVSPVVFDVLFLFGIFNSCVNPIVYGLSMFKKTTARPTLSLIEFSSPFLTRSERRSASVNSRISRTYSHVSLSTRRSWQPSSESTTSSRPNALNNQHSHSASHLMTGLPCRYGVSRQVRGGSQQKYLHPNTTTSPGPSTRSPKEASWHLKRATLTHAHSSPSLLTSMEKSERPNRRQSEALAVPTLPTICLTPPSESRNTAFFSAALCLLHDEYANSPTQPPVADIPYQPTAVNQQGDQHYLSRRLSVALPMIPPTPDTPGSWETSSWQFVGKISCSLCRYDSSSPSASYPDRGECVTGRPQVNARRCISLGNTDSKVNPKITERLNAEVVQKMRNSVQETKISEYDLSFEVASFRPPLHRLSVQESCVQSSGRMASGASIVGSEGSIQSSLESPLHSPASSVTSIDFRHRLSLCSFREGRIPSDLHTALRPKHYSDDNLKLSCRKRKAKRRVSFKTVHFLAEPTVSDSSVPSEGSNSSTPDDHQEPTRARNTSNHRTVSADIKWHVGCQLTRKTSSTLP

>Branchiostoma_floridae_CRZR2__ACN79527_1 _FJ426561.1_(Formerly_GnRHR4)

CCACCATGAGCCCATCCCAGTCTCCCCGCACCGAGAACGACAGCGCCTTCTTTCGGGAGGACGTGTTCCTGAAGGATTTTCTGAACGAGAGCTCCGATCCAGAGCTGATCAATGAGACCTCCAGAAACAATAAGACAATGGACAGGGATCTGAACCTGCCCGAGTTCACCACATGGACCCTGACAAAGATCGTGACCATCTGCATCCTGTTCGTGATCGCCGCCATCGGCAATCTGTTTATGGCCCGCGCCACACTGAGGCTGCGGAGAAGGAGCGGCATCTACCTGCTGCTGCTGCACCTGTCCGTGGGCGAGCTGCTGGTGACCTGCATCACAATGCCTTCTGAGGCCATCTGGGCATACACCGTGAGCTGGTGGGCAGGCGACACAATGTGCAGAATCGTGAAGTATGGCCAGATGCTGGGCCTGTACCTGTCCACCTATATCACAGTGTGCATCTCTCTGGACAGATGCGTGGCCATCGCATTCCCCCTGAAGAAGGGACAGGCACCTGAGAGGGCACGGTCCATGGTCATCGTGAGCTGGGCCCTGAGCCCTATCTTCTGCATCCCACAGGCCGTGATCTTTCACGTGGAGGTGCACCAGTACGTGCCATCTTTCCACCAGTGCGTGACCTACAACTTTTATAGCGCCGAGTGGCAGGAGGACCTGTACAATATGCTGGTGTTTGTGGTCATGTATCCCGCCCCTATGGTCATCATGGTGGCCTGCTACGTGTGCATCTTCGTGTCCCTGTTTAGACACTGGAGGGGCACAAACAATCTGGAGACCGGCAATAAGACAGGCCAGAGAGAGAGGCTGTTCTCTAAGGCAAAGGTGCGCACCCTGCAGATGGCAGCCGGCATCCTGACCACCTTCTTCGTGTGCTGGACACCCTTTTACTGCGTGATGATGTGGCACCTGTTCTTTCAGCACGAGTACCCTATCAACCAGATCATCTTCGACGTGCTGTATCCATTTGGCGTGAGCAACGCCTGCGTGAATCCCGTGGTGTATGGCAAGTCCGTGGTGACCAGAAAGCCTGGCAAGTCTTTCCTGGTGAATTGGTACCAGGCCTTTCTGGAGCCAGAGGAGTATGCCCGCAAGCTGGACTCTACCAAGGCCGCCGCCTGTACAAGGCTGTCTAGCCTGCGCCGGAGCGAGCAGAGGGATAGGATGAACTCCCTGACCCCCACAGTGTACGTGGAGGTGAGCAGAAGGAATTCCTCTAGAATCGTGACCCAGGATCAGCTGAGCGGCTCCAGGGTGTGA

>Protein_Branchiostoma_floridae_CRZR2__ACN79527_1_XP_035667678_1_FJ426561.1_(Formerly_GnRHR4)

MSPSQSPRTENDSAFFREDVFLKDFLNESSDPELINETSRNNKTMDRDLNLPEFTTWTLTKIVTICILFVIAAIGNLFMARATLRLRRRSGIYLLLLHLSVGELLVTCITMPSEAIWAYTVSWWAGDTMCRIVKYGQMLGLYLSTYITVCISLDRCVAIAFPLKKGQAPERARSMVIVSWALSPIFCIPQAVIFHVEVHQYVPSFHQCVTYNFYSAEWQEDLYNMLVFVVMYPAPMVIMVACYVCIFVSLFRHWRGTNNLETGNKTGQRERLFSKAKVRTLQMAAGILTTFFVCWTPFYCVMMWHLFFQHEYPINQIIFDVLYPFGVSNACVNPVVYGKSVVTRKPGKSFLVNWYQAFLEPEEYARKLDSTKAAACTRLSSLRRSEQRDRMNSLTPTVYVEVSRRNSSRIVTQDQLSGSRV

>Branchiostoma_floridae_CRZR3_XP_035665381.1_GnRHR_like5

CCACCATGACTCCAACTTCTGTGTCCATAGACACCACAGACGCGGACAACGTGACGAATTCGACCATTTCGTCGTATCCGGAGGTGCCAACCTTCAGCGACCACCACCTGATGAAAGTCGTCATCACTTCGTTCCTGTTTCTGTGGGCCGCCGTAGGGAACGTGTCGGTGTTGGCGGTGATGCAGCGGTCGCGCTCCCGGAGCAGGACTCCTCTCCACTCCCTGGTGTTCCAGCTCACGGTCGCAGACTCCCTGGTCACCTTCATCACCATGCCGTCTGAGGCCGCCTGGGCCGCCACCGTGTCGTGGAAGGCTGGAGAAGTCATGTGCAGAGTCATCAAGTTTCTGCAGGTTATTGGACTGTACCAGTCGACGTTTATCACGGTGGCGATAAGTCTGGACCGGGCGGTCGCCGTGCTGTTCCCCTTCAGTAAGAGCGGAGCTCCACATCGCACCAAGGTCATGATCATCATCTGCTGGGTCCTCAGCACAGTCTTCAGCTTCCCGCAGGCGATCATTTTTCACATCGAGCGACTTGTCCCGGGGTTTGAGCAATGCGTCGACTTCAACTTCTACGGAGCCAAGTGGCAGAAACAGCTGTACAACACCGCGTCGTTCGTCCTCATCTACCCGCTGCCTCTCGTCATCATCATCACGGCTTACGTCTGCATCATTCTCCGTATCACCAAGAACACAAGGACCAAAGGTAACGCCTTTTCCGGATCACAGCACGTTCGGTCTGAAATGGACGCCAGAAGGGAGCAAGTCTTTACCAGGGCAAAGATCCGGACATTGTGGATGACGGTGGGCATTGTCACTGCCTTCGTGGTGTGCTGGACGCCATTTTATGTCCACCTGTTCTGGGTAAACTACTTCGACACCAGCCATGTCTCCAGGATGCTGACCGACCTGCTGTATCTGTTCGGGATGTCCAACGCTTGTGTCAACCCCATGGTGTACGGCCTCAGCACGTTTCTCCGCGCGCCTCCCAACTCCAACAAGACCCGCTCACAAACTCTGCACGTCCTCTACACTCTCAAGAACCACACCGACAACAGCTGCGTGAGTAGGAGTCCATCAAGACGCATTAATAACGTGGACAAGACCGTTTTAGTGCATTCTACCGTTAGTCAGGAATCAGCCACAAATGGCGAACACATGCACAGCAAATACAGAGAAACTTCCTGCTGA
>Protein_Branchiostoma_floridae_CRZR3_XP_035665381.1_GnRHR_like5

MTPTSVSIDTTDADNVTNSTISSYPEVPTFSDHHLMKVVITSFLFLWAAVGNVSVLAVMQRSRSRSRTPLHSLVFQLTVADSLVTFITMPSEAAWAATVSWKAGEVMCRVIKFLQVIGLYQSTFITVAISLDRAVAVLFPFSKSGAPHRTKVMIIICWVLSTVFSFPQAIIFHIERLVPGFEQCVDFNFYGAKWQKQLYNTASFVLIYPLPLVIIITAYVCIILRITKNTRTKGNAFSGSQHVRSEMDARREQVFTRAKIRTLWMTVGIVTAFVVCWTPFYVHLFWVNYFDTSHVSRMLTDLLYLFGMSNACVNPMVYGLSTFLRAPPNSNKTRSQTLHVLYTLKNHTDNSCVSRSPSRRINNVDKTVLVHSTVSQESATNGEHMHSKYRETSC
